# DISCO+QR: Rooting Species Trees in the Presence of GDL and ILS

**DOI:** 10.1101/2023.01.02.522492

**Authors:** James Willson, Yasamin Tabatabaee, Baqiao Liu, Tandy Warnow

## Abstract

Genes evolve under processes such as gene duplication and loss (GDL), so that gene family trees are multi-copy, as well as incomplete lineage sorting (ILS); both processes produce gene trees that differ from the species tree. The estimation of species trees from sets of gene family trees is challenging, and the estimation of rooted species trees presents additional analytical challenges. Two of the methods developed for this problem are STRIDE (Emms and Kelly, MBE 2017), which roots species trees by considering GDL events, and Quintet Rooting (Tabatabaee et al., ISMB 2022 and Bioinformatics 2022), which roots species trees by considering ILS. We present DISCO+QR, a new method for rooting species trees in the presence of both GDL and ILS. DISCO+QR, operates by taking the input gene family trees and decomposing them into single-copy trees using DISCO (Willson et al., Systematic Biology 2022) and then roots the given species tree using the information in the single-copy gene trees using Quintet Rooting (QR). We show that the relative accuracy of STRIDE and DISCO+QR depend on properties of the dataset (number of species, genes, rate of gene duplication, degree of ILS, and gene tree estimation error), and that each provides advantages over the other under some conditions. Availability: DISCO and QR are available in GitHub. The supplementary materials are available at http://tandy.cs.illinois.edu/discoqr-suppl.pdf.

## 1 Introduction

The estimation of rooted species trees is a basic step in many biological research problems, including the detection of selection, understanding adaptation, the timing of evolutionary events, etc. Over the last several years (and especially the last decade) there has been remarkable progress in developing methods that have strong statistical guarantees and high accuracy for estimating species trees, typically from multi-locus datasets, but these methods only produce unrooted trees [16, 27, 7, 18].

Several methods have been developed and are in wide use for rooting estimated species trees, including using an outgroup species (i.e., a species that is not in the clade defined by the other species) to root the species tree and midpoint rooting (i.e., rooting the species tree at the midpoint of the longest leaf-to-leaf path). While both of these methods are popular, neither is completely reliable. For example, outgroup selection is challenging since: (i) if the putative outgroup is a true outgroup but very distantly related, finding the edge to which the outgroup species attaches can be difficult (due to accumulation of mutations) and (ii) some putative outgroups may be in fact not outgroups (a challenge that can occur when the root of the phylogeny is a rapid radiation [24]). For these, as well as other reasons [20], outgroup selection is challenging and unreliable in practice. The problem with midpoint rooting is that evolutionary changes are not always clocklike [2, 19], and when the deviation from the clock is sufficient, midpoint rooting can be inaccurate. Thus, these two standard approaches to rooting species trees are unreliable.

In the last few years, new methods have been developed that take a statistical approach to rooting species trees from multi-gene datasets based on models of gene evolution within species trees. For example, STRIDE [8] is a method for rooting species trees from multi-copy gene family trees that result from gene duplications and losses (GDL). Another method that has been developed is Quintet Rooting (QR) [25], which is a method for rooting species trees from single-copy gene trees that can differ from the species tree due to incomplete lineage sorting (ILS).

In this study, we present a new technique for rooting species trees that can address both GDL and ILS. Our new method is a combination of QR, which can only work with single-copy gene trees and is based on ILS alone, and a previous method, DISCO [29], for breaking gene family trees into single-copy gene trees. The combination of these two methods, DISCO+QR, operates by using DISCO to break each gene family tree into single-copy gene trees and then applies QR to the resultant single-copy gene trees. This surprisingly simple approach is fast (polynomial time and fast in practice). Our simulation study evaluating DISCO+QR in comparison to STRIDE shows that each has a range of conditions where it is more accurate than the other, with STRIDE generally more accurate when ILS is low and GDL rates are high, and DISCO+QR more accurate when ILS is high.

## 2 Background

### 2.1 Estimating unrooted species trees under GDL+ILS

When genes evolve under processes that include duplications and losses, then their phylogenies have multiple copies of each species, so that the gene trees are referred to as gene family trees (or MUL-trees). Many methods have been developed for estimating species trees from gene family trees, with gene tree parsimony approaches (i.e., minimizing the total number of duplications and losses, and weighted versions of these criteria) [4, 28] perhaps the most well-known. Other approaches include variations of supertree methods to allow for MUL-tree inputs; examples of such methods include MulRF [5] and FastMulRFS [17]. Yet, none of these methods has been proven statistically consistent for estimating the species tree.

#### 2.1.1 Quartet-based methods

The first results for statistical consistency and identifiability under any GDL model were provided in [12], who proved that for every four species, the most frequently observed quartet tree (among the gene family trees) would be topologically identical to the species tree. This result established that properly designed quartet-based methods would be statistically consistent for species tree estimation in the presence of GDL. This theorem was later extended to show quartet-based methods, when properly designed, would be statistically consistent under a joint model for GDL and ILS (the DLCOAL model [22]) in [15].

Subsequent to these studies that established statistical identifiability using quartet trees (and hence potential statistical consistency of methods based on quartet trees), quartet-based methods have been developed for species tree estimation in the presence of GDL and ILS, and some of these have been proven statistically consistent. Specifically, ASTRAL-one and ASTRAL-multi are two ways of running the ASTRAL species tree estimation method (which is also based on quartet trees) that are established statistically consistent under the DLCOAL model. ASTRAL-Pro [31] is a new variant of ASTRAL specifically designed to address GDL; several simulation studies have shown it is more accurate than ASTRAL-multi and ASTRAL-one, but it has not yet been proven statistically consistent.

#### 2.1.2 ASTRID-DISCO

Another approach for estimating species trees from gene family trees that has very good accuracy, comparable to ASTRAL-Pro (sometimes more accurate) but also faster, is ASTRID-DISCO, which we now describe.

ASTRID [27] is a polynomial time method for estimating unrooted species trees from unrooted gene trees that is statistically consistent under the multispecies coalescent model; thus, ASTRID addresses species tree estimation when ILS is present, but does not address GDL. ASTRID operates by computing the matrix of average internode distances between species, and then runs distance-based methods (e.g., FastME [11] with balanced minimum evolution) to produce the species tree.

DISCO is a method for decomposing a gene family tree (which can contain multiple copies of each species) into a set of single copy gene trees. DISCO operates using two steps. In the first step, it “roots-and-tags” the gene family tree, which means that it finds a root location that implies the smallest total number of duplication and loss events, and uses that rooting to identify the nodes that are duplication events (this is the “tagging” step); this is the same technique used in ASTRAL-Pro. The second step then decomposes the tree into single copy trees at the duplication events, seeking to keep at least one large subtree. The two steps together thus decompose a single gene family tree into a set of single-copy gene trees. As shown in [29], if ASTRAL-Pro correctly roots and tags the gene family tree, then the single-copy gene trees produced by DISCO contain only orthologs.

In ASTRID-DISCO, each gene family tree in the input set of gene family trees is split into single copy trees using DISCO [29], and then ASTRID is applied to the set of single copy trees to estimate the species tree. As shown in [29], ASTRID-DISCO was generally more accurate than the other leading species tree estimation methods, including ASTRAL-Pro [31], under a wide range of model conditions. Moreover, ASTRID-DISCO is very fast, since both ASTRID and DISCO are low-degree polynomial time; and in particular, ASTRID-DISCO is faster than ASTRAL-Pro. Thus, ASTRID-DISCO matches or improves on the accuracy and speed of ASTRAL-Pro and other methods for species tree estimation.

### 2.2 Rooting species trees using STRIDE

STRIDE is a method designed to root species trees given a set of unrooted multi-copy gene family trees, assuming that all gene tree heterogeneity is due to gene duplication and loss. It determines a set of “strongly supported” duplication events given an unrooted species tree and the unrooted gene family trees, then uses this set of duplications to find the best rooting for the species tree. STRIDE outputs both a species tree—rooted on the edge it judges to be the best—and a file listing all of the maximum parsimony roots (roots that minimize the number of duplication events needed to explain them) and the probability that each edge could be the root.

In the study in [8], STRIDE was applied to a collection of biological datasets from different groups of species and three previously published simulated datasets, where two of the simulated datasets contained both GDL and ILS. Most of these datasets had very large numbers of gene family trees (ranging from 8, 392 to 28, 356 for the biological datasets and on average ≈ 7, 200 for the simulated datasets) and varying numbers of species, from 5 to 47.

STRIDE had very good performance on these datasets, and was able to find the correct root on 12 out of 15 datasets. Although STRIDE was not directly compared to any other rooting method, its gene duplication prediction accuracy was compared to two reconciliation methods, Notung [6] and DLCpar_search [30] and was shown to outperform them.

### 2.3 Rooting species trees using Quintet Rooting (QR)

QR is a method for rooting species trees given a set of unrooted single-copy gene trees, assuming that all gene tree heterogeneity is due to ILS. QR is based on the proof of identifiability of rooted species trees with at least 5 taxa from unrooted gene trees under the MSC provided by Allman, Degnan and Rhodes (ADR) [1]. Specifically, ADR establish a set of inequalities and invariants that must hold among the probabilities of the true gene trees under each 5-taxon model species tree, and prove that these inequalities and invariants are sufficient to identify the rooted species tree topology on 5 taxa.

QR scores each possible rooting of a given unrooted species tree based on the probability distribution that it computes over a selected set of quintets in the gene trees. It operates by selecting a set of quintets of taxa from the leafset of the input tree (referred to as “quintet sampling”), and in a preprocessing step, it computes a cost for each possible rooting of these quintet trees. The score of each rooted tree is then computed by summing up the costs over all its induced rooted quintets, and the tree with the minimum score is returned as the best possible rooting. Similar to STRIDE, QR also returns an additional file containing a score and a ranking for all possible rootings of the input tree (each corresponding to the root being located on one edge in the unrooted tree).

QR can be used with different quintet sampling methods that differ in terms of their runtime and accuracy. Two sampling methods proposed in [25] are “exhaustive” and “linear encoding (LE)” samplings, where the first computes a cost for all 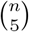 quintets of taxa, but the second looks at a carefully chosen set of *O*(*n*) quintets (where *n* is the number of species) using an encoding that maps each edge in the unrooted tree to a single quintet. A previous study in [25] showed that the exhaustive mode of QR is generally more accurate than LE, as it is based on more information; however, it is significantly slower. QR is polynomial-time when used with both sampling methods, and the detailed runtime analysis with each sampling method is provided in Section 3.

In [25], QR was compared with four other rooting methods, midpoint [10], Minimum Ancestor Deviation (MAD) [26], Minimum Variance Rooting (MinVar) [13] and RootDigger [3], and was shown to have better or comparable accuracy to other rooting methods on a set of simulated datasets and 5-taxon subsets of an avian biological dataset. The experimental study in [25] explored rooting both true and estimated species trees given true or estimated single-copy gene trees as input, when ILS was the only source of discordance, and QR had very good relative accuracy on these datasets except when gene tree estimation error was very high.

## 3 New method: DISCO+QR

DISCO+QR is a new approach for rooting species trees given a set of gene family trees. Operationally it is very simple: Given the input gene family trees, DISCO is used to split each gene family tree into single-copy trees. Then, QR is used to root the input species tree with respect to the single-copy trees. The idea behind DISCO+QR is that if DISCO is successful, then each single-copy tree contains only orthologs, and any remaining discordance between the resulting single-copy gene trees and the species trees is assumed due to ILS and potentially gene tree estimation error. Under these circumstances, the species tree can be rooted with high accuracy if there is enough ILS, so that the single copy gene trees can identify the root location.

### Theorem 1.

*The time complexity of DISCO+QR is O*(*N*(|*Q*| + *n*)), *where Q is the set of sampled quintets in QR, N is the total number of leaves in all multi-copy gene family trees and n is the number of taxa in the species tree. Hence, when using the linear encoding, DISCO+QR uses O*(*nN*) *time*.

*Proof*. We divide the DISCO+QR pipeline into three steps:

1. Run DISCO to decompose the gene-family trees into single-copy gene trees.
2. QR Preprocessing: Compute the cost for each of the seven possible rootings of an unrooted quintet tree induced on each quintet in set *Q*.
3. QR Scoring: Compute the score for all 2*n* − 3 different rootings of the input unrooted species tree.

In Proposition S1.1, we prove that the runtime of the DISCO decomposition step is *O*(*Nn*). In Proposition S1.2, we prove that the preprocessing step of QR is done in *O*(*k*(|*Q*| + *n*)) time, where *k* is the number of single-copy gene trees given as input to QR. Note that *k* ≤ *N* since DISCO can output at most *N* single-copy gene trees, and therefore the runtime of QR preprocessing is bounded by *O*(*N*(|*Q*| + *n*)). In Proposition S1.3 we prove that the scoring step of QR can be done in *O*(*n* + |*Q*|), where *n* is the number of taxa in the species tree. The total runtime of QR is the sum of the three parts, which is *O*(*N*(|*Q*| + *n*) + *n*). In particular, when QR is used with linear encoding (LE) sampling, the total running time of DISCO+QR reduces to *O*(*Nn*).

The proofs of Propositions S1.1, S1.2 and S1.3 are provided in the supplementary materials, Section S1.

## 4 Experimental Study

### 4.1 Overview

We use a collection of simulated datasets (some new and others from prior studies) to evaluate DISCO+QR in comparison to STRIDE. Our datasets vary in terms of levels of ILS (low and moderate) and GDL (six different levels), and have varying numbers of genes. Our model conditions use gene trees estimated using maximum likelihood and so vary in terms of estimation error. We ran four experiments, with Experiments 1–3 examining conditions with 20 species and two levels of ILS, and Experiment 4 examined larger numbers of species (up to 1000). In each experiment we evaluated accuracy on the root location when given a species tree estimated using ASTRID-DISCO (described above).

We report error in the root location using average normalized clade distance (nCD), which is an extension of Robinson-Foulds (RF) [23] distance for rooted trees and can be computed using the following formula

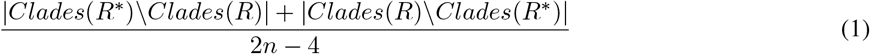

where *R*^*^ and *R* are true and estimated rooted species trees respectively and *Clades*(*t*) is the set of all clades of a rooted tree *t*. The clade distance is also a measure of distance to the root, and is equal to twice the root distance (distance of the estimated root position to the true root) when rooting the true species tree topology [25].

For full details of the experimental study, see the supplementary materials, available at http://tandy.cs.illinois.edu/discoqr-suppl.pdf. All software and datasets are freely available (see Appendix 8).

### 4.2 Datasets

We used simulated data containing GDL, ILS, and gene tree estimation error (GTEE) from [29], as well as new data simulated for this study.

#### 4.2.1 Data Simulation

We simulated the new data for this study using a pipeline similar to [17, 31, 29]: First the true species tree and the true gene trees are simulated with SimPhy [14]. We use INDELible [9] to evolve sequences down the true gene trees and then estimate gene trees from these sequences with the maximum likelihood method, FastTree2 [21].

The duplication rates and loss rates are controlled directly through SimPhy, while we control the level of ILS using the haploid effective population size. We also set several options to ensure that the evolution diverges from a strict molecular clock (i.e., branch rate heterogeneity modifiers). Our new datasets contain 21 species (20 species and an additional outgroup taxon), 1000 gene trees per replicate, and varies the duplication rates under two levels of ILS; all model conditions also contain two levels of gene tree estimation error (estimated from 100bp and 50bp sequences). The ILS level is measured in terms of average distance (AD), which is the average normalized Robinson-Foulds (RF) distance between the locus trees (which are true gene family trees with GDL but before modification by ILS) and true gene family trees (which reflect both ILS and GDL). More details, including the commands used, are in the Supplementary Materials.

## 5 Results

### 5.1 Experiment 1: Varying the duplication rate

Our first experiment compared differing duplication rates at two levels of ILS (Fig. 1), while holding the number of gene trees to 1000 and the amount of estimation error to moderate (about 40% MGTE). The clearest trend is that STRIDE is greatly affected by the duplication rate; at moderate to extremely high duplication rates (1 × 10^−11^ to 1 × 10^−9^), it performs well; however, as the duplication rate lowers (1 × 10^−12^ and 1 × 10^−13^), its performance degrades. On the other hand, DISCO+QR seems largely unaffected by the duplication rates; instead, its performance is mostly impacted by the ILS levels. At the greatest level of ILS, it performs almost as well as STRIDE, even outperforming it at low levels of GDL, while with little ILS, its accuracy degrades. STRIDE does worse at high rates of ILS compared to low rates of ILS, but the differences are very small for all but the lowest duplication rate.

**Figure 1:**
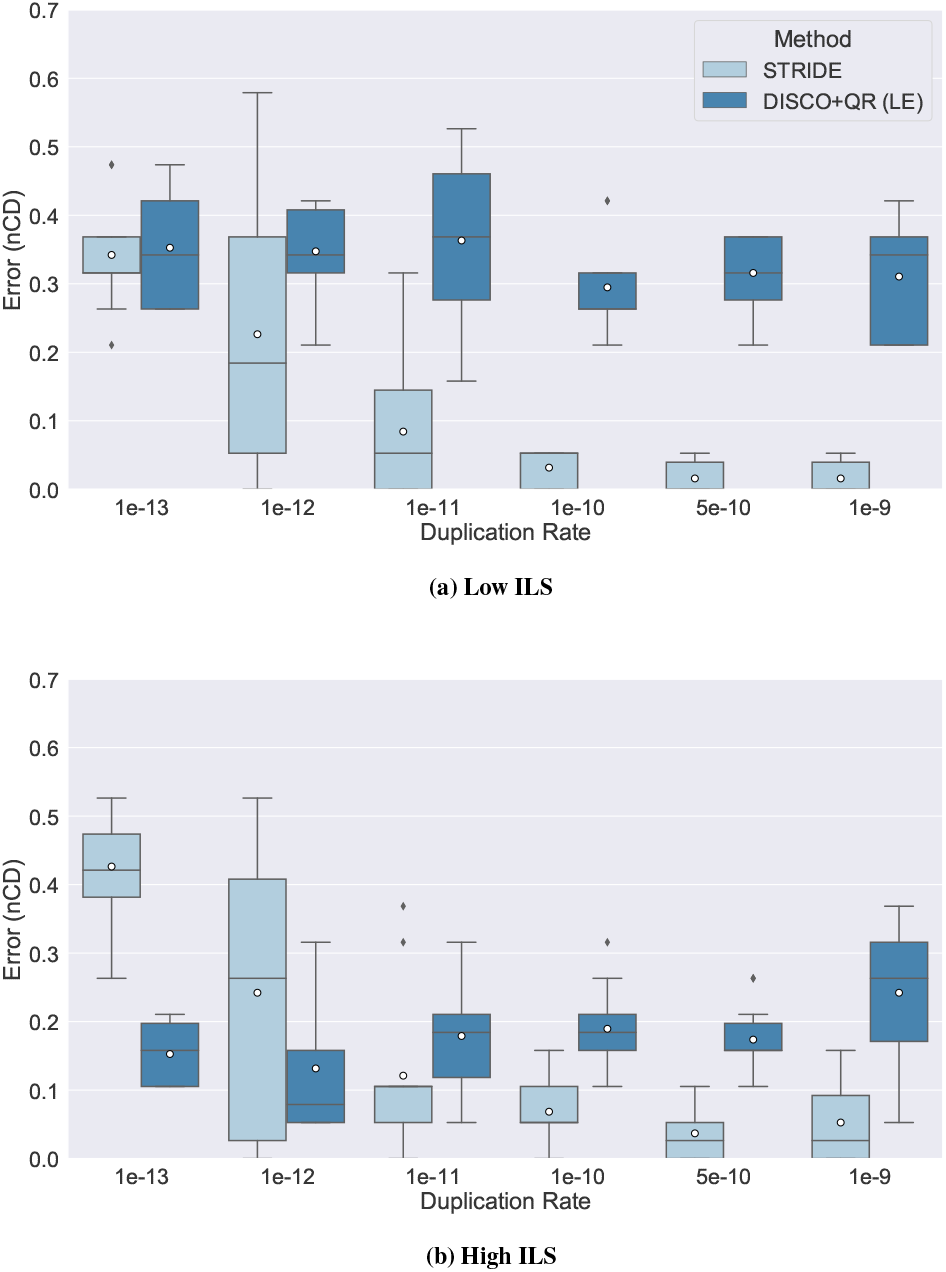
Results on 21-species datasets, showing impact of varying the duplication rate on rooting error (nCD) of STRIDE and DISCO+QR, given at two levels of ILS; low ILS in subfigure (a) (AD ≈ 20%) and high ILS in subfigure (b) (AD ≈ 64%). The species tree was estimated with ASTRID-DISCO. The simulated model conditions contain 1000 genes, 10 replicates, a loss rate equal to the duplication rate, and ≈ 40% MGTE. Note: circles are means and bars are medians.

### 5.2 Experiment 2: Varying gene tree estimation error

In Experiment 2, we evaluated the effect of gene tree estimation error (GTEE) on both methods (Fig. 2). Notably, we keep the duplication rate fixed at 1 × 10^−12^, where STRIDE outperforms DISCO+QR under the condition with low ILS and DISCO+QR outperforms STRIDE with high ILS. These trends continue under all levels of gene tree estimation error. The impact of ILS on STRIDE, however, is much clearer under conditions with very low gene tree estimation error (the two lowest conditions examined); under these low GTEE conditions, STRIDE’s performance is much worse when ILS is high than when it is low.

**Figure 2:**
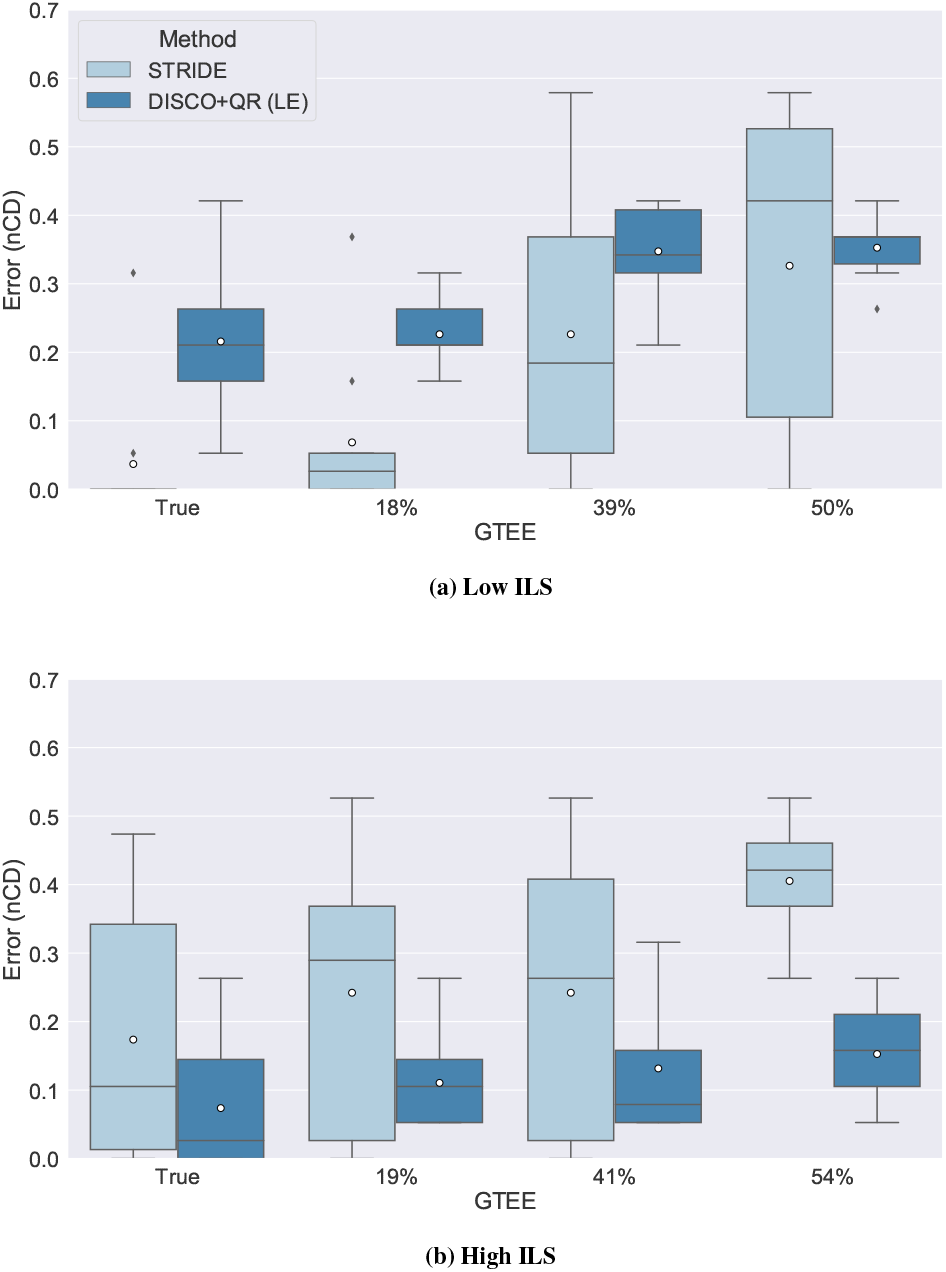
Results on 21-species datasets, showing impact of varying the amount of gene tree estimation error on the rooting error (nCD) of STRIDE and DISCO+QR, given at two levels of ILS; low ILS in subfigure (a) (AD ≈ 20%) and high ILS in subfigure (b) (AD ≈ 64%). The species tree was estimated with ASTRID-DISCO. The simulated model conditions contain 1000 genes, 10 replicates, a duplication rate of 1 *×* 10^−12^ and an equal loss rate, and AD ≈ 20%. Note: circles are means and bars are medians.

### 5.3 Experiment 3: Varying the number of genes

In this experiment we varied the number of genes (Fig. 3). Again we fix the duplication rate at 1 *×* 10^−12^ and examine both levels of ILS. As expected, both methods become more accurate as the number of genes increases, but the relative accuracy remains the same (with DISCO+QR more accurate than STRIDE for high ILS and STRIDE more accurate than DISCO+QR for low ILS). Under the high ILS condition, the number of gene trees have a larger impact on the accuracy of both methods than under the low ILS condition (where both methods accuracy seems largely unaffected until 1000 genes).

**Figure 3:**
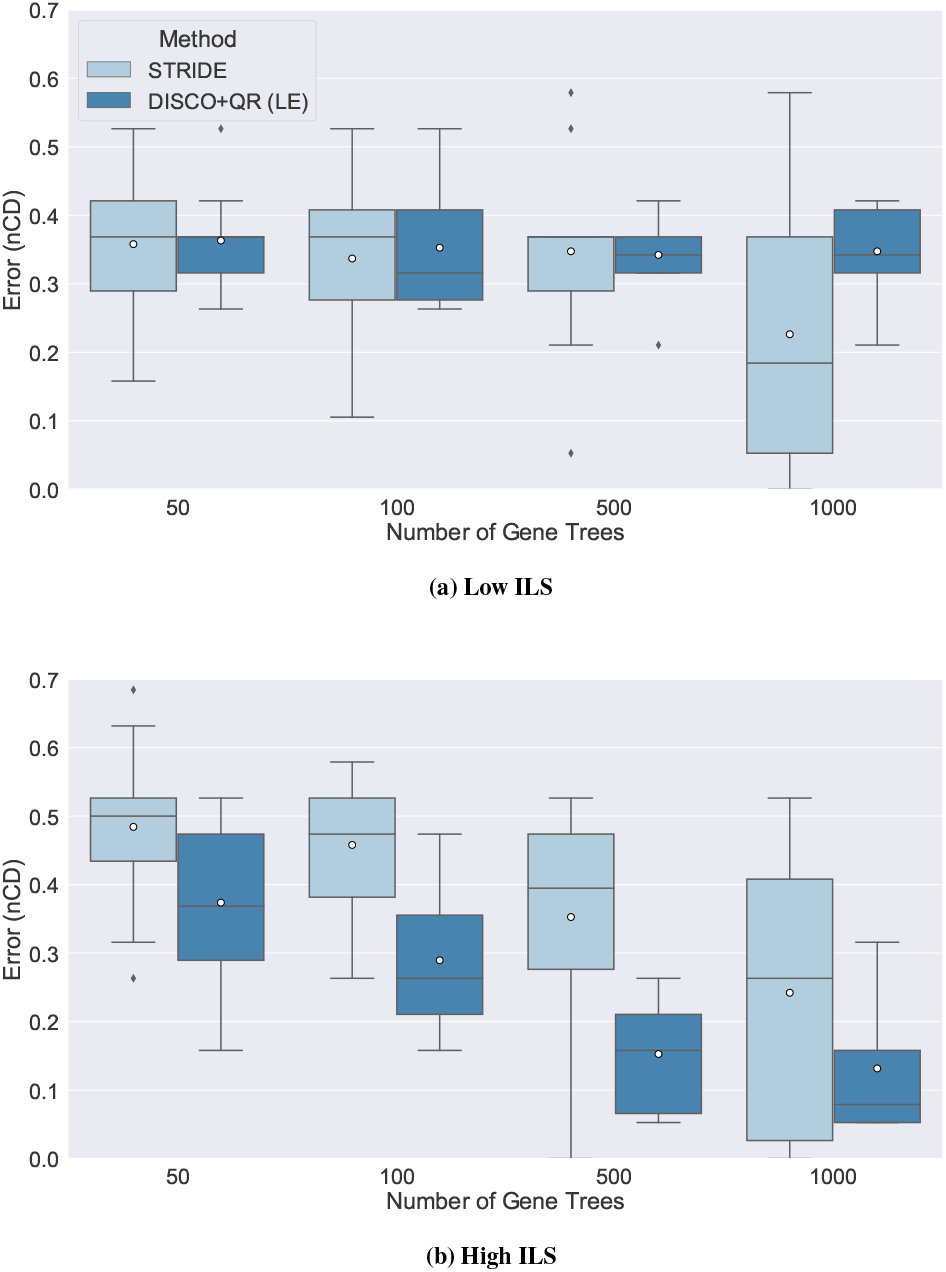
Results on 21-species datasets, showing impact of varying the number of gene trees on the rooting error (nCD) of STRIDE and DISCO+QR, given at two levels of ILS; low ILS in subfigure (a) (AD ≈ 20%) and high ILS in subfigure (b) (AD ≈ 64%). The species tree was estimated with ASTRID-DISCO. The simulated model conditions contain 10 replicates, a duplication rate of 1 *×* 10^−12^ and an equal loss rate, AD ≈ 20%, and ≈ 40% MGTE. Note: circles are means and bars are medians.

### 5.4 Experiment 4: Examining larger numbers of species

We now examine trends on larger numbers of species (up to 1000).

#### 5.4.1 Moderate duplication rate and 20–100 species

Figure 4 examines conditions with 1000 genes and moderate duplication rate (1 *×* 10^−12^), equal loss rate, moderate gene tree estimation error, two levels of ILS (low and moderately high), while varying the number of species from 20 to 100. The first observation is that in general error rates decrease with the number of species (although the benefit to DISCO+QR under high ILS is relatively small). The second observation is that DISCO+QR continues to be more accurate than STRIDE for all numbers of species for the high ILS condition, but the gap between the two methods decreases with the number of species. The third observation is that while STRIDE is more accurate than DISCO+QR at 20 species and low ILS, the two are essentially tied at 50 species and then DISCO+QR is more accurate at 100 species (but only by a small amount).

**Figure 4:**
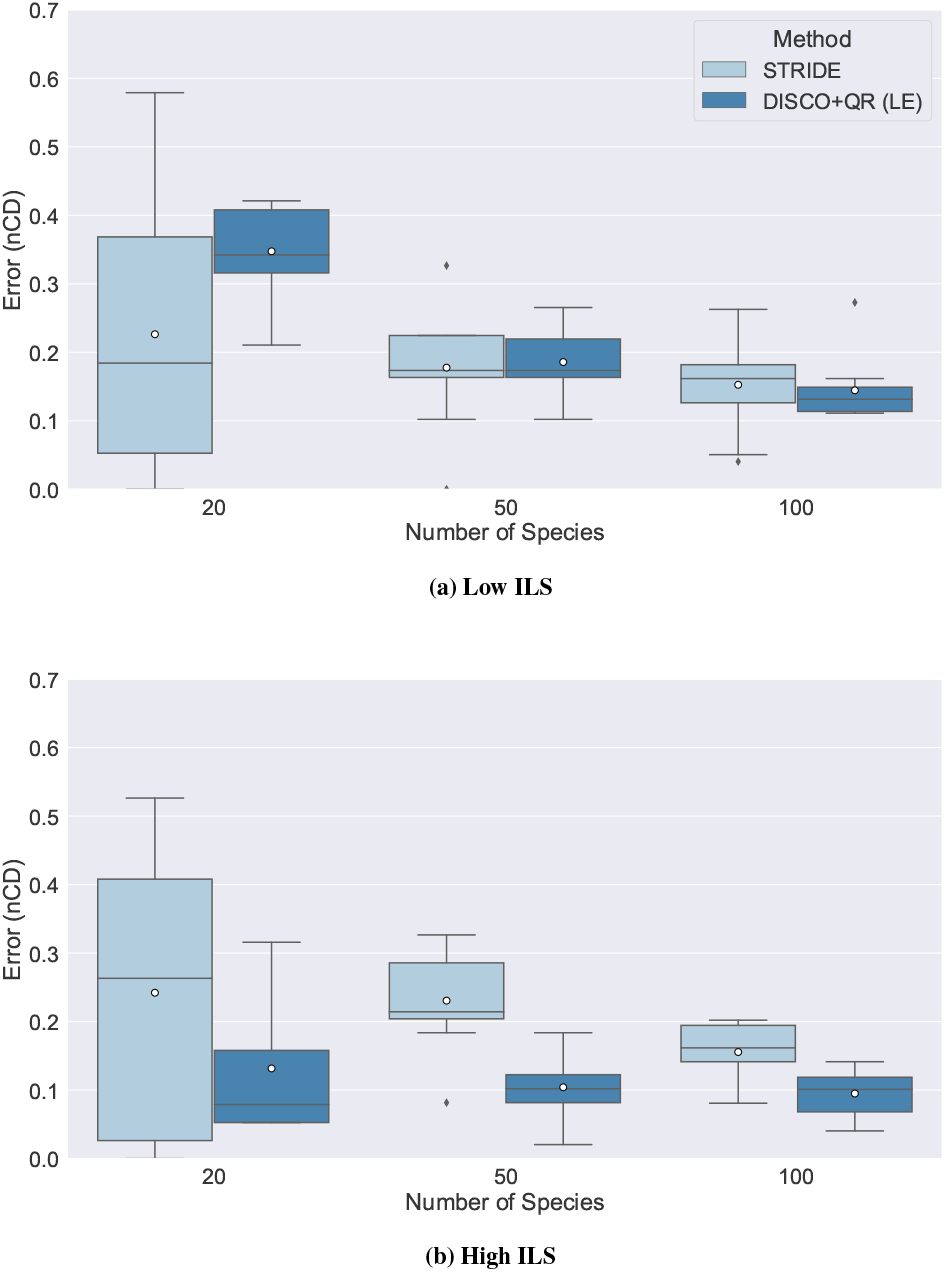
Impact of varying the number of species on the rooting error (nCD) of STRIDE and DISCO+QR, given moderate duplication rates. (a) low ILS (AD ≈ 20%), (b) high ILS (AD ≈ 64%). The species tree was estimated with ASTRID-DISCO. The simulated model conditions contain 1000 genes, 10 replicates, a moderate duplication rate of 1 *×* 10^−12^ and an equal loss rate, and ≈ 40% MGTE. Note: circles are means and bars are medians.

Thus, although our experiments on 20 species consistently showed an advantage of STRIDE over DISCO+QR when ILS was low, this advantage is no longer shown on larger numbers of species, where DISCO+QR was tied with STRIDE at 50 species and then more accurate than STRIDE on 100 species. However, this specific experiment was only under a moderate duplication rate, leaving open the question of what would happen when the duplication rate is higher.

#### 5.4.2 High duplication rates, low ILS, and 100 species

The next experiment evaluated methods with l00 species and high duplication rates. Figure 5 shows results as we vary the number of genes from 100 to 1000, for three high duplication rates (varying from 1 *×* 10^−10^ to 1 *×* 10^−9^). The following trends are clear. First, at 1000 genes, STRIDE is very accurate, with mean nCD ranging from 0.033 to 0.057, while DISCO+QR has higher mean nCD that range from 0.112 to 0.133. For 500 genes, STRIDE is still highly accurate, with mean nCD between 0.045 and 0.065 at all tested duplication rates, and DISCO+QR is again less accurate but still reasonable (mean nCD between 0.126 and 0.151). At 100 genes, the error increases for both methods (mean nCD for STRIDE between 0.116 and 0.277, and between 0.147 and 0.318 for DISCO+QR), so that while STRIDE maintains an advantage over DISCO+QR, the gap between the two methods has decreased.

**Figure 5:**
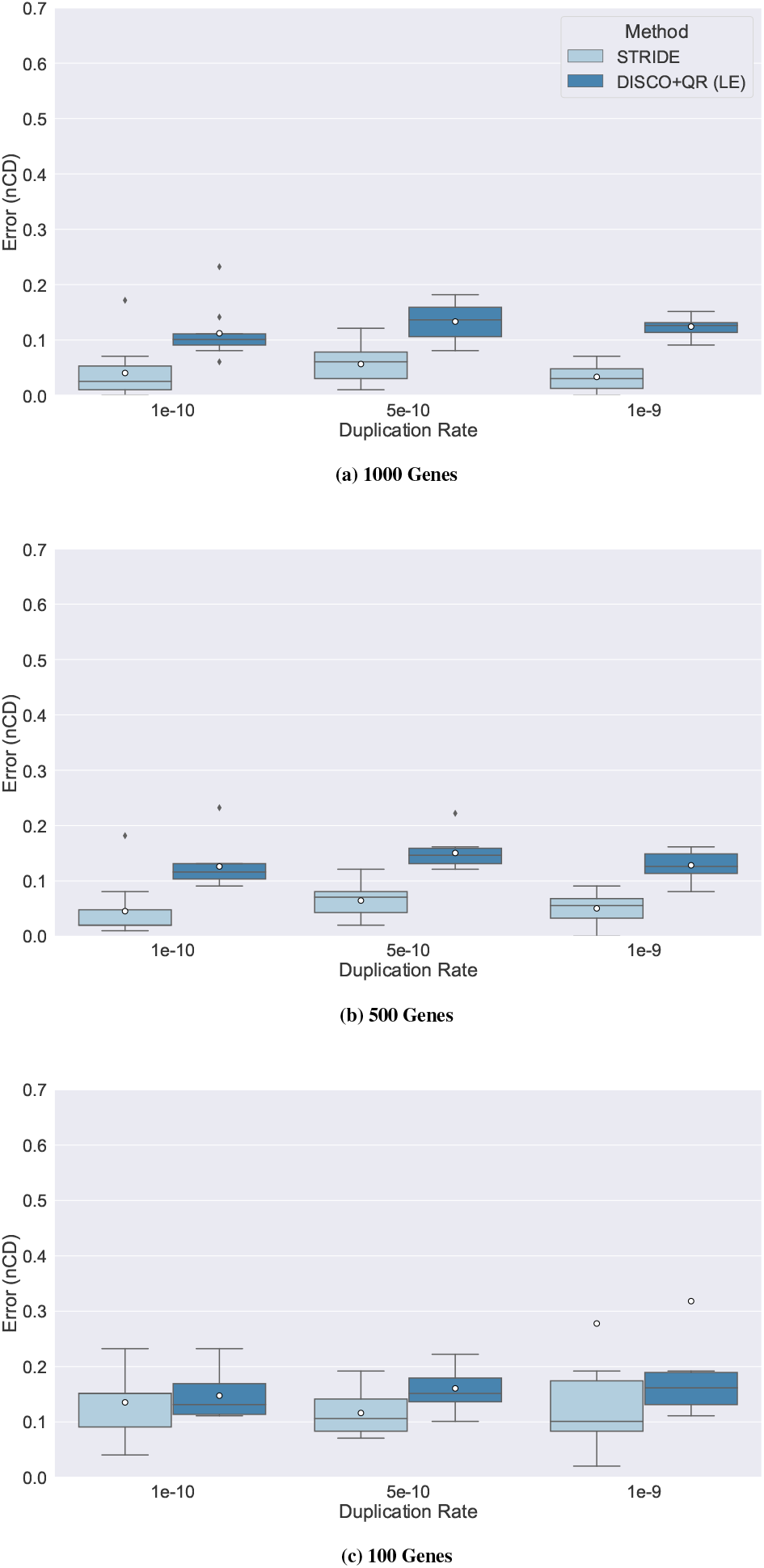
Results on 101-species datasets, evaluating impact of varying number of genes under three high duplication rates and low ILS on the rooting error (nCD) of STRIDE and DISCO+QR on simulated datasets with varying numbers of genes. (a) 1000 genes, (b) 500 genes, (c) 100 genes. The species tree was estimated with ASTRID-DISCO. The simulated model conditions contain 10 replicates, a loss rate equal to the duplication rate, AD ≈ 20%, and ≈ 40% MGTE. Note: circles are means and bars are medians.

#### 5.4.3 High duplication rates, low ILS, and 1000 species

Results on 1000 species, 100–1000 genes, and high duplication rates are shown in Figure 6. At 1000 genes, both STRIDE and DISCO+QR have excellent accuracy, with mean nCD of 0.054 and 0.066, respectively. Moreover, with lower numbers of genes, error increases for both methods but only modestly, so that at 500 genes the mean nCDs are still very low (0.06 for STRIDE and 0.07 for DISCO+QR). Interestingly, with only 100 genes, DISCO+QR (mean nCD of 0.081) has an accuracy advantage over STRIDE (mean nCD of 0.262); the explanation is that STRIDE has some outliers with very high (1.0) nCD values.

**Figure 6:**
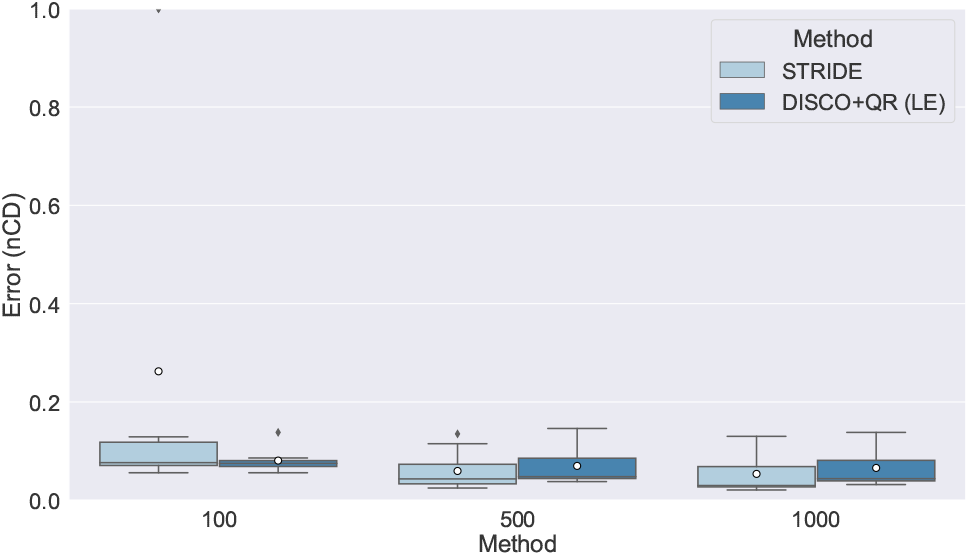
Results on 1001-species datasets, with varying numbers of genes. The species tree was estimated with ASTRID-DISCO. The simulated model conditions contain 10 replicates, a duplication rate of 5 *×* 10^−10^ and an equal loss rate, AD ≈ 20%, and ≈ 40% MGTE. Note: circles are means and bars are medians.

Overall, therefore, for the 1000-species condition we explored (a low ILS condition with high duplication rates), the two methods are extremely close for 500 and 1000 genes (never off by more than 0.012 in nCD), but with an advantage to DISCO+QR for the 100-gene case. This is a surprising contrast to the datasets with fewer species.

## 6 Discussion

This study evaluated STRIDE and DISCO+QR under a range of model conditions with varying ILS, GDL, number of genes, number of species, and gene tree estimation error. For both methods, accuracy increased with number of genes and decreases in gene tree estimation error; this is consistent with many other studies. For conditions with only 20 species, in general STRIDE had an advantage over DISCO+QR when there was low ILS and DISCO+QR had an advantage over STRIDE under high ILS. The relative performance of the methods was also impacted by duplication rates, with low duplication rates favoring DISCO+QR and high duplicaiton rates favoring STRIDE. As the number of species increased from 20 to as high as 1000, the trends changed, with DISCO+QR beginning to show advantages over STRIDE even when ILS was low and duplication rates were higher. The results on 1000 species are especially interesting, since on that model condition the two methods are extremely close on 500 or more genes, but with an advantage to DISCO+QR for 100 genes. The model condition also has low ILS and high GDL, both of which would seem to favor STRIDE based on the conditions with smaller numbers of species. This finding, although limited to just a small part of the parameter space, suggests some potential advantage to DISCO+QR on very large datasets that merits further investigation.

## 7 Conclusion

This study presented DISCO+QR, a new method for rooting species trees, given a set of gene family trees that evolve under both gene duplication and loss (GDL) and incomplete lineage sorting (ILS). DISCO+QR is low degree polynomial time and fast enough to be used on large datasets with 1000 species and 1000 genes. DISCO+QR is typically more accurate than STRIDE under high ILS conditions, and typically less accurate under low ILS conditions; however, the experimental study also suggests that relative accuracy will depend on a combination of factors, including number of species, number of genes, levels of gene tree estimation error, and GDL rates.

This study suggests several directions for future work. For example, given the fact that STRIDE and DISCO+QR each have advantages over the other depending on these dataset properties, it seems likely that taking input from both methods may provide better techniques for rooting species trees. We also note that modifications to the DISCO+QR pipeline (e.g., changes to the QR optimization problem) may improve its accuracy. This study mainly examined small numbers of species, yet DISCO+QR clearly improves significantly in accuracy on larger numbers of species; hence, further study of conditions with more species is necessary. Future work could also evaluate the accuracy of alternative methods, such as midpoint rooting, for rooting an estimated species tree. One challenge in such an approach is scalability to large datasets (specifically with large numbers of species and loci), due to the computational effort in estimating branch lengths and other numeric parameters on a large species tree. Additionally, future work should also examine results on biological datasets, where the true root is known or considered highly reliable, as was performed in [8].

## 8 Data and Software Availability

Links to the data used in this study are provided at http://tandy.cs.illinois.edu/datasets.html. DISCO is available on GitHub at https://github.com/JSdoubleL/DISCO and QR at https://github.com/ytabatabaee/Quintet-Rooting. The supplementary materials are available at http://tandy.cs.illinois.edu/discoqr-suppl.pdf.

## 10 Author contributions statement

TW: Conceptualization, Supervision, Project administration, Funding Acquisition, Methodology, Writing - Original draft preparation, Writing - Reviewing and Editing. JW: Methodology, Investigation, Software, Visualization, Writing - Original draft preparation, Writing - Reviewing and Editing. YT: Methodology, Investigation, Software, Writing- Original draft preparation, Writing - Reviewing and Editing. BL: Methodology, Investigation, Software, Writing - Original draft preparation, Writing - Reviewing and Editing.

## 11 Acknowledgments

This work was supported by the Department of Computer Science at UIUC and by the Grainger Foundation.

