## supplement for "DISCO+QR: Rooting Species Trees in the Presence of GDL and ILS"

**S3 Additional Information about Datasets** 6

**S4 Additional Results** 8

### S1 Proof of Runtime Complexity of DISCO+QR

Given an unrooted tree $T$, we denote a rooted version of $T$ by $R_e$ where $R_e$ is $T$ rooted on edge $e$. If the rooted edge is not of significance, we omit $e$ and simply write $R$.

**Proposition S1.1.** *DISCO decomposes the input gene family trees into single copy trees in $O(Nn)$ time, where $N$ is the total number of leaves in the gene family trees and $n$ is the number of taxa (species labels) appearing in the gene trees.*

*Proof.* The DISCO algorithm can be separated into three steps for each input gene-family tree $G_i$. The first step, rooting (finding the most parsimonious root), can be done in $O(n|L_i|)$ time, where $|L_i|$ is the number of leaves in $G_i$. This is briefly mentioned in the original algorithm description in [7] ("we used memoization to reduce the time to quadratic"). The second step, tagging, can be again done in $O(n|L_i|)$ time, which is clear from its algorithm in ASTRAL-Pro (Algorithm 1, [7]). The third step, decomposition, can be done in $O(|L_i|)$ time, which is self-evident from its pseudo-code in the DISCO paper (Algorithm 1, [6]). The total running time of the DISCO decomposition step for all input gene family trees is thus $\sum_i O(|L_i|n) = O(nN)$ □

The next two propositions give the runtime of the two steps of QR: preprocessing and scoring. First, we bring a detailed description of the QR algorithm:

**Details of QR Algorithm.** QR has a cost function $Cost(R_e|_q, \vec{u}_q)$ that given $\vec{u}_q$ (i.e., $\vec{u}_q$ a 15-dimensional vector), the empirical distribution of gene tree unrooted quintet topologies induced on $q$ and a rooted quintet topology $R_e|_q$ (denoting $R$ rooted on $e$ induced on $q$), calculates the "cost" of observing the distribution $\vec{u}_q$ assuming $R_e|_q$ is the model rooted quintet topology. Given the set of sampled quintets $Q$, $Score(e)$ is defined as $\sum_{q\in Q} Cost(R_e|_q, \vec{u}_q)$. Our goal is to find the edge $e$ that minimizes $Score(e)$. Now we give the scoring algorithm.

1. Preprocess $T$. Build an empty list $I(v) \subseteq Q$ at each internal node $v$ of $T$. For each $q \in Q$, consider $T|_q$. We extract the internal nodes $x, y, z$ of $T|_q$, where $x, y, z$ are also internal nodes of $T$. Add $q$ to $I(x)$, $I(y)$, and $I(z)$. In effect, $I(v)$ for any internal node $v$ of $T$ is defined by the quintets inside $Q$ such that the homeomorphic subtrees of $T$ induced on these quintets have $v$ as an internal node.

2. Fix an arbitrary edge $e$. Calculate $Score(e)$ by iterating over the quintets in $Q$.

3. Consider an edge $f$ adjacent to $e$ connected by node $v$. We conceptually split $Q$ into two sets. The first set $Q_\delta$ is defined by those $q \in Q$ s.t. $R_e|_q \neq R_f|_q$ in topology, i.e., those quintets that have rooted topologies that differ when rerooting $T$ from $e$ to $f$. The second set $Q'$ is $Q - Q_\delta$ (quintets that don't differ in rooted topology when rerooting from $e$ to $f$). Observe that for any $q' \in Q'$, $R_e|_{q'} = R_f|_{q'}$ in topology, implying $Cost(R_e|_{q'}, \vec{u}_{q'}) = Cost(R_f|_{q'}, \vec{u}_{q'})$. Therefore,
$$Score(f) - Score(e) = \sum_{q\in Q_\delta} Cost(R_f|_q, \vec{u}_q) - Cost(R_e|_q, \vec{u}_q)$$

   Further observe that $Q_\delta = I(v)$, since by construction, for any $q \in I(v)$, $e$ and $f$ are separated by an internal node of $T|_q$ that is $v$, hence $R_e|_q$ and $R_f|_q$ are different in rooted topology. Also, any $q \notin I(v)$ will be in $Q'$ because $e$ and $f$ are located on the same edge of $T|_q$ (where an edge in $T|_q$ is a path in $T$).

4. Finally, recall that we already calculated $Score(e)$ for some arbitrary $e$. We proceed by traversing all edges of $T$ starting from $e$, and when visiting an edge $f$, we derive $Score(f)$ using the above-mentioned update strategy. The above strategy will be called at most twice per internal node $v$ because each $v$ has bounded degree three.

We now bring the runtime complexity analysis of preprocessing and scoring steps.

**Proposition S1.2.** *The preprocessing step of QR that computes the cost of each possible rooting of an unrooted quintet can be done in $O(k(|Q| + n))$, where $k$ is the number of input single-copy gene trees and $Q$ is the set of sampled quintets in QR.*

*Proof.* Recall that the cost of a rooted quintet tree on taxon-set $q$ is defined with respect to an empirical distribution of the unrooted quintet topologies of $q$ observed in the gene trees denoted by $\vec{u}_q$. Therefore we first obtain this $\vec{u}_q$, after which the cost for a given rooted quintet tree can be computed in $O(1)$ time.

For each single-copy gene tree $G$, we first equip $G$ with an LCA data structure (see Lemma S1.4) in $O(n)$ time, after which the unrooted quintet topology of any $q$ can be queried on $G$ in $O(1)$ time (by Lemma S1.5). As such, each $\vec{u}_q$ can be computed in $O(k)$ time, and all $\vec{u}_q$s where $q \in Q$ can be computed in $O(|Q|k)$ time. Adding in the time to initialize the LCA data structures results in a total running time of $O((|Q| + n)k)$.

Once the $\vec{u}_q$s are computed, one can iterate over all seven possible rooted topologies of $T|_q$, where $T|_q$ is the unrooted species tree induced on $q$, to compute the costs against $\vec{u}_q$. Since there are a constant number of topologies per $q$, our total running time is unaffected and remains $O((|Q| + n)k)$. □

**Proposition S1.3.** *The scoring step of QR can be done in $O(n+|Q|)$ time for an unrooted species tree $T$ with $n$ taxa, where $Q$ is the set of quintets sampled.*

*Proof.* Given that $R_e|_q$ can be queried in $O(1)$ time according to Lemma S1.5, each $\vec{u}_q$ already calculated in the QR preprocessing step, and that $Cost$ is a constant time function, $Score(e)$ can be derived in $O(|Q|)$ time for arbitrary $e$ by simply iterating over all quintets in set $Q$. Now we compute the runtime for each step of the QR scoring algorithm:

1. Preprocessing $T$ takes $O(n)$ time according to Lemma S1.4, and computing $I(v)$ for all internal nodes of $T$ takes $O(|Q|)$ time, as each $q \in Q$ is analyzed in constant time. Therefore, this step takes $O(n + |Q|)$ time.

2. Calculating $Score(e)$ for one fixed edge $e$ can be simply done by iterating over all the quintets in set $Q$. This takes $O(|Q|)$ time.

3. For an edge $f$ connected to $e$ by node $v$, updating $Score(e)$ to $Score(f)$ is done in $O(|I(v)|)$ time, since $Q_\delta = I(v)$ and $Q_\delta$ contains quintets $q$ such that $R_e|_q$ and $R_f|_q$ are different in rooted topology and only the cost of these quintets should be updated in $Score(f)$, which can be done in constant time per quintet.

4. Each internal node of $T$ will be traversed at most twice in the final step, and hence the scores of all edges can be computed in at most $\sum_v 2 \cdot (1 + O(|I(v)|))$ time.

In this way, we obtained scores for all edges of $T$ in at most $\sum_v 2 \cdot (1 + O(|I(v)|)) = O(n + |Q|)$ time, where $v$ ranges over all internal nodes of $T$. Therefore, the total runtime of the scoring step of QR is $O(n + |Q|)$. □

**Lemma S1.4.** *Given rooted tree $R$ with $l$ leaves, after $O(l)$ preprocessing time, the following queries can be completed in $O(1)$ time for arbitrary nodes $u, v$ of $R$.*

1. *$lca(u, v)$, the least common ancestor of $u$ and $v$*
2. *$depth(u)$, the distance (number of edges) between $u$ and the root. If $depth(u) > depth(v)$, $u$ is said to be deeper (or lower) than $v$.*
3. *$d(u, v)$, the number of edges between $u$ and $v$*
4. *$ancestor(u, v)$, which decides if $v$ is an ancestor of $u$*

*Proof.* These are well known. (1) is due to [2]. (2) is trivial (preprocess by DFS from the root) and is frequently a byproduct of the preprocessing used for (1). For (3), $d(u, v) = depth(u) + depth(v) - 2 \cdot depth(lca(u, v))$ and hence can be queried in constant time. For (4), $u$ is an ancestor of $v$ if $lca(u, v) = u$ and hence can be queried in constant time. □

We simply refer to this preprocessing of Lemma S1.4 as equipping $R$ with the LCA data structure.

**Lemma S1.5.** *Given rooted tree $R$ (with $T$ as its unrooted topology) equipped with the LCA data structure, the following queries can be completed in $O(1)$ time given five taxa $q = (q_1, q_2, q_3, q_4, q_5)$ and $e$ an edge in $R$:*

1. *$R|_q$, the homeomorphic (rooted) subtree of $R$ induced on $q$*
2. *$T|_q$, the homeomorphic subtree of $T$ induced on $q$*
3. *$R_e|_q$, first reroot $R$ on $e$, then consider the rooted quintet topology of $R_e$ induced on $q$*

*Proof.* We start with (1). Consider the set $S$ of nodes in $R$ inductively defined as follows. First, $q_i \in S$ where $1 \leq i \leq 5$. Second, if $u, v \in S$, then $lca(u, v) \in S$. Observe that $S$ is constant-size by construction and contains exactly all the nodes of $R|_q$. Construct $S$ in constant time, and for any $u, v \in S$, if $u$ is an ancestor of $v$, put a directed edge from $u$ to $v$, forming a graph $H$ on $S$. Compute the transitive reduction on $H$, which results in $R|_q$. Notice that $R|_q$ is of constant size.

For (2), since $R|_q$ is of constant size, we can use brute force to extract the two bipartitions of the unrooted topology of $R|_q$ in constant time, which defines $T|_q$.

For (3), first observe that given two nodes $x, y$ in $R$ and an edge $e = (u, v)$ in $T$, it can now be decided in constant time if $e$ lies on the path between $x$ and $y$. A node $z$ is between $x, y$ iff $d(x, z) + d(z, y) = d(x, y)$. Therefore, if $u$ and $v$ both lie between $x, y$, then $e$ lies between $x, y$. First compute $R|_q$. Then, among all edges of $R|_q$, we need to find the edge $x, y$ s.t. $e$ is located on the path between $x$ and $y$ on $T$, which as we just established can be done in constant time per edge, and we have a constant number of edges. Then reroot $R|_q$ on this edge $x, y$ on which $e$ lies, obtaining our defined $R_e|_q$. □

### S2 Software Commands and Version Numbers

#### S2.1 DISCO+QR

DISCO and Quintet Rooting (QR) are available in open source form in github (as described below). Therefore, DISCO+QR can be easily run by a simple pipeline.

#### S2.2 STRIDE

We ran STRIDE (version 1.0.0) rooting software which is available at https://github.com/davidemms/STRIDE using the following command

```
python2 stride.py -s dash -S <species_tree> -o <output_prefix> -d <gene_tree_dir>
```

where `<gene_tree_dir>` is a directory containing gene family trees in separate files.

The `-o` option is enabled in this forked version https://github.com/JSdoubleL/STRIDE.

#### S2.3 Quintet Rooting (QR)

We ran Quintet Rooting (QR) (version 1.2.2) available at https://github.com/ytabatabaee/Quintet-Rooting with the following command:

```
python3 quintet_rooting.py -t <species-topology.tre> -g <input-genes.tre>
-o <output-tree.tre> -sm LE -rs 0
```

The `-sm LE` option specifies the quintet sampling method as “Linear Encoding”.

#### S2.4 DISCO

We ran DISCO (version 1.3) available at https://github.com/JSdoubleL/DISCO for decomposing multi-copy gene-family trees into single-copy gene trees. We used the following command:

```
python3 disco.py -i <gene_trees> -o <output_file> -d _
```

#### S2.5 Rooting (Clade Distance) Error, MGTE, AD

Mean gene tree estimation error (MGTE) and average distance (AD) between the model species tree and true gene trees were calculated using a script written by Erin. K. Molloy for computing normalized Robinson-Foulds (RF) distance, available at

https://github.com/ekmolloy/fastmulrfs/blob/master/python-tools/compare_two_trees.py.

We measured the rooting error using average normalized clade distance (nCD) which is an extension of RF distance for rooted trees. We used the script for computing average normalized clade distance available at https://github.com/ytabatabaee/Quintet-Rooting/blob/main/scripts/clade_distance.py which is extended and modified from the script written by Erin. K. Molloy for computing RF distance for unrooted trees.

#### S3 Additional Information about Datasets

To create the new simulated data for this study, we used SimPhy [3] with the following command:

```
simphy -sl f:20 -rs 10 -rl f:1000 -rg 1 -sb f:0.000000005 \
    -sd f:0 -st ln:21.25,0.2 -so f:1 -si f:1 -sp f:$population_size \
    -su ln:-21.9,0.1 -hh f:1 -hs ln:1.5,1 \
    -hl ln:1.551533,0.6931472 -hg ln:1.5,1 -cs 8472 -v 3 \
    -o $model_condition_name -ot 0 -op 1 -lb f:$dup_rate \
    -ld f:$loss_rate -lt f:0
```

To simulate gene tree estimation error, we needed to generate sequence data. We used INDELible [1] with GTR parameters taken from the FastMulRFS study [4]. We also used scripts from that study (available here: https://databank.illinois.edu/datasets/IDB-5721322) to run INDELible.

```
python set_indelible_params.py \
    -n 1000 \
    -f 113.48869 69.02545 78.66144 99.83793 \
    -r 12.776722 20.869581 5.647810 9.863668 30.679899 3.199725 \
    -a -0.470703916 0.348667224 \
    -l $sequence_length \
    -o indelible-parameters.csv

python run_indelible.py \
    -x indelible \
    -s 1 \
    -e 1000 \
    -p indelible-parameters.csv \
    -t g_true.trees \
    -o $out_dir
```

Finally, we generated trees from the simulated sequences using FastTree2 [5], with the follwing command:

```
fasttree -nt -gtr $alignment_file > $estimated_tree
```

| Dup Rate | # Taxa | AD | MGTE (500bp) | MGTE (100bp) | MGTE (50bp) | Avg. # Leaves |
|---|---|---|---|---|---|---|
| | | | Low ILS – Equal Loss | | | |
| $1 \times 10^{-13}$ | 21 | 0.184 | – | 0.379 | 0.490 | 21.0 |
| $1 \times 10^{-12}$ | 21 | 0.221 | 0.184 | 0.415 | 0.525 | 21.0 |
| $1 \times 10^{-12}$ | 51 | 0.209 | – | 0.421 | – | 51.0 |
| $1 \times 10^{-12}$ | 101 | 0.251 | – | 0.459 | – | 101.2 |
| $1 \times 10^{-11}$ | 21 | 0.251 | – | 0.422 | 0.534 | 21.3 |
| $1 \times 10^{-10}$ | 21 | 0.190 | – | 0.368 | 0.478 | 24.1 |
| $5 \times 10^{-10}$ | 21 | 0.158 | – | 0.342 | 0.452 | 35.8 |
| $1 \times 10^{-9}$ | 21 | 0.156 | – | 0.337 | 0.441 | 52.1 |
| | | | High ILS – Equal Loss | | | |
| $1 \times 10^{-13}$ | 21 | 0.693 | – | 0.422 | 0.554 | 21.0 |
| $1 \times 10^{-12}$ | 21 | 0.674 | 0.192 | 0.426 | 0.557 | 21.0 |
| $1 \times 10^{-12}$ | 51 | 0.757 | – | 0.458 | – | 51.2 |
| $1 \times 10^{-12}$ | 101 | 0.784 | – | 0.482 | – | 101.2 |
| $1 \times 10^{-11}$ | 21 | 0.670 | – | 0.410 | 0.532 | 21.3 |
| $1 \times 10^{-10}$ | 21 | 0.645 | – | 0.390 | 0.509 | 24.2 |
| $5 \times 10^{-10}$ | 21 | 0.545 | – | 0.395 | 0.517 | 36.2 |
| $1 \times 10^{-9}$ | 21 | 0.442 | – | 0.383 | 0.507 | 47.9 |
| | | | Low ILS – No Loss | | | |
| $1 \times 10^{-13}$ | 21 | 0.210 | – | 0.380 | 0.482 | 21.0 |
| $1 \times 10^{-12}$ | 21 | 0.200 | – | 0.391 | 0.500 | 21.1 |
| $1 \times 10^{-11}$ | 21 | 0.173 | – | 0.401 | 0.511 | 21.7 |
| $1 \times 10^{-10}$ | 21 | 0.215 | – | 0.388 | 0.499 | 30.0 |
| $5 \times 10^{-10}$ | 21 | 0.191 | – | 0.416 | 0.541 | 158.2 |
| $1 \times 10^{-9}$ | 21 | 0.195 | – | 0.409 | 0.550 | 693.9 |
| | | | High ILS – No Loss | | | |
| $1 \times 10^{-13}$ | 21 | 0.666 | – | 0.416 | 0.545 | 21.0 |
| $1 \times 10^{-12}$ | 21 | 0.674 | – | 0.423 | 0.547 | 21.1 |
| $1 \times 10^{-11}$ | 21 | 0.674 | – | 0.417 | 0.542 | 21.8 |
| $1 \times 10^{-10}$ | 21 | 0.628 | – | 0.411 | 0.538 | 30.8 |
| $5 \times 10^{-10}$ | 21 | 0.589 | – | 0.405 | 0.550 | 147.3 |
| $1 \times 10^{-9}$ | 21 | 0.558 | – | 0.461 | 0.604 | 776.4 |

Table S1: Statistics for simulated model conditions containing 10 replicates with 1000 gene trees each. High ILS indicates the model conditions with the high simulated haploid effective population size, while low indicates the reverse.

#### S4 Additional Results

This section has tables with means and medians for the results (given in figures) from the main paper.

| Model Condition | Method | Mean | Median |
|---|---|---|---|
| Figure 1(a): Low ILS | | | |
| $1 \times 10^{-13}$ | DISCO+QR (LE) | 0.353 | 0.342 |
| $1 \times 10^{-12}$ | | 0.347 | 0.342 |
| $1 \times 10^{-11}$ | | 0.363 | 0.368 |
| $1 \times 10^{-10}$ | | 0.295 | 0.263 |
| $5 \times 10^{-10}$ | | 0.316 | 0.316 |
| $1 \times 10^{-9}$ | | 0.311 | 0.342 |
| $1 \times 10^{-13}$ | STRIDE | 0.339 | 0.316 |
| $1 \times 10^{-12}$ | | 0.210 | 0.184 |
| $1 \times 10^{-11}$ | | 0.103 | 0.000 |
| $1 \times 10^{-10}$ | | 0.017 | 0.000 |
| $5 \times 10^{-10}$ | | 0.016 | 0.000 |
| ] $1 \times 10^{-9}$ | | 0.012 | 0.000 |
| Figure 1(b): High ILS | | | |
| $1 \times 10^{-13}$ | DISCO+QR (LE) | 0.153 | 0.158 |
| $1 \times 10^{-12}$ | | 0.132 | 0.079 |
| $1 \times 10^{-11}$ | | 0.179 | 0.184 |
| $1 \times 10^{-10}$ | | 0.189 | 0.263 |
| $5 \times 10^{-10}$ | | 0.174 | 0.158 |
| $1 \times 10^{-9}$ | | 0.242 | 0.263 |
| $1 \times 10^{-13}$ | STRIDE | 0.400 | 0.421 |
| $1 \times 10^{-12}$ | | 0.226 | 0.263 |
| $1 \times 10^{-11}$ | | 0.142 | 0.078 |
| $1 \times 10^{-10}$ | | 0.062 | 0.026 |
| $5 \times 10^{-10}$ | | 0.045 | 0.000 |
| $1 \times 10^{-9}$ | | 0.055 | 0.000 |

Table S2: Means and medians for all model conditions from Figure 1. Impact of varying the duplication rate on rooting error (nCD) of STRIDE and DISCO+QR, given at two levels of ILS; low ILS in subfigure (a) (AD $\approx$ 20%) and high ILS in subfigure (b) (AD $\approx$ 64%). The species tree was estimated with ASTRID-DISCO. The simulated model conditions contain 21 taxa, 1000 genes, 10 replicates, a loss rate equal to the duplication rate, and $\approx$ 40% MGTE.

| Model Condition | Method | Mean | Median |
|---|---|---|---|
| Figure 2(a): Low ILS | | | |
| True | DISCO+QR (LE) | 0.216 | 0.211 |
| 18% | | 0.226 | 0.211 |
| 39% | | 0.351 | 0.342 |
| 50% | | 0.353 | 0.368 |
| True | STRIDE | 0.037 | 0.000 |
| 18% | | 0.036 | 0.026 |
| 39% | | 0.226 | 0.184 |
| 50% | | 0.326 | 0.421 |
| Figure 2(b): High ILS | | | |
| True | DISCO+QR (LE) | 0.074 | 0.026 |
| 18% | | 0.111 | 0.105 |
| 39% | | 0.132 | 0.079 |
| 50% | | 0.153 | 0.158 |
| True | STRIDE | 0.174 | 0.105 |
| 18% | | 0.242 | 0.289 |
| 39% | | 0.242 | 0.263 |
| 50% | | 0.405 | 0.421 |

Table S3: Means and medians for all model conditions from Figure 2. Impact of varying the amount of gene tree estimation error on the rooting error (nCD) of STRIDE and DISCO+QR, given at two levels of ILS; low ILS in subfigure (a) (AD $\approx$ 20%) and high ILS in subfigure (b) (AD $\approx$ 64%). The species tree was estimated with ASTRID-DISCO. The simulated model conditions contain 21 taxa, 1000 genes, 10 replicates, a duplication rate of $1 \times 10^{-12}$ and an equal loss rate, and AD $\approx$ 20%.

| Model Condition | Method | Mean | Median |
|---|---|---|---|
| Figure 3(a): Low ILS | | | |
| 50 | DISCO+QR (LE) | 0.363 | 0.368 |
| 100 | | 0.353 | 0.316 |
| 500 | | 0.342 | 0.342 |
| 1000 | | 0.347 | 0.342 |
| 50 | STRIDE | 0.358 | 0.368 |
| 100 | | 0.337 | 0.368 |
| 500 | | 0.347 | 0.368 |
| 1000 | | 0.226 | 0.184 |
| Figure 3(b): High ILS | | | |
| 50 | DISCO+QR (LE) | 0.374 | 0.368 |
| 100 | | 0.289 | 0.263 |
| 500 | | 0.152 | 0.158 |
| 1000 | | 0.132 | 0.079 |
| 50 | STRIDE | 0.484 | 0.500 |
| 100 | | 0.458 | 0.474 |
| 500 | | 0.353 | 0.395 |
| 1000 | | 0.242 | 0.263 |

Table S4: Mean and median for all model conditions from Figure 3. Impact of varying the number of gene trees on the rooting error (nCD) of STRIDE and DISCO+QR, given at two levels of ILS; low ILS in subfigure (a) (AD $\approx$ 20%) and high ILS in subfigure (b) (AD $\approx$ 64%). The species tree was estimated with ASTRID-DISCO. The simulated model conditions contain 21 taxa, 10 replicates, a duplication rate of $1 \times 10^{-12}$ and an equal loss rate, AD $\approx$ 20%, and $\approx$ 40% MGTE.

| Model Condition | Method | Mean | Median |
|---|---|---|---|
| Figure 4(a): Low ILS | | | |
| 20 | | 0.347 | 0.342 |
| 50 | DISCO+QR (LE) | 0.186 | 0.173 |
| 100 | | 0.144 | 0.131 |
| 20 | | 0.226 | 0.184 |
| 50 | STRIDE | 0.178 | 0.173 |
| 100 | | 0.153 | 0.162 |
| Figure 4(b): High ILS | | | |
| 20 | | 0.132 | 0.079 |
| 50 | DISCO+QR (LE) | 0.104 | 0.102 |
| 100 | | 0.095 | 0.101 |
| 20 | | 0.242 | 0.263 |
| 50 | STRIDE | 0.231 | 0.214 |
| 100 | | 0.155 | 0.162 |

Table S5: Means and medians for all model conditions from Figure 4. Impact of varying the number of species on the rooting error (nCD) of STRIDE and DISCO+QR, given moderate duplication rates. (a) low ILS (AD $\approx$ 20%), (b) high ILS (AD $\approx$ 64%). The species tree was estimated with ASTRID-DISCO. The simulated model conditions contain 1000 genes, 10 replicates, a moderate duplication rate of $1 \times 10^{-12}$ and an equal loss rate, and $\approx$ 40% MGTE.

| Model Condition | Method | Mean | Median |
|---|---|---|---|
| Figure 5(a): 1000 genes | | | |
| $1 \times 10^{-10}$ | | 0.112 | 0.101 |
| $5 \times 10^{-10}$ | DISCO+QR (LE) | 0.133 | 0.136 |
| $1 \times 10^{-9}$ | | 0.124 | 0.126 |
| $1 \times 10^{-10}$ | | 0.040 | 0.025 |
| $5 \times 10^{-10}$ | STRIDE | 0.057 | 0.061 |
| $1 \times 10^{-9}$ | | 0.033 | 0.030 |
| Figure 5(b): 500 genes | | | |
| $1 \times 10^{-10}$ | | 0.126 | 0.116 |
| $5 \times 10^{-10}$ | DISCO+QR (LE) | 0.151 | 0.146 |
| $1 \times 10^{-9}$ | | 0.128 | 0.126 |
| $1 \times 10^{-10}$ | | 0.045 | 0.020 |
| $5 \times 10^{-10}$ | STRIDE | 0.065 | 0.071 |
| $1 \times 10^{-9}$ | | 0.050 | 0.056 |
| Figure 5(c): 100 genes | | | |
| $1 \times 10^{-10}$ | | 0.147 | 0.131 |
| $5 \times 10^{-10}$ | DISCO+QR (LE) | 0.161 | 0.152 |
| $1 \times 10^{-9}$ | | 0.318 | 0.162 |
| $1 \times 10^{-10}$ | | 0.135 | 0.152 |
| $5 \times 10^{-10}$ | STRIDE | 0.116 | 0.106 |
| $1 \times 10^{-9}$ | | 0.277 | 0.101 |

Table S6: Mean and median for all model conditions from Figure 5. Impact of varying number of genes under three high duplication rates and low ILS on the rooting error (nCD) of STRIDE and DISCO+QR on simulated datasets with 100 taxa (and one outgroup taxon) and varying numbers of genes. (a) 1000 genes, (b) 500 genes, (c) 100 genes. The species tree was estimated with ASTRID-DISCO. The simulated model conditions contain 10 replicates, a loss rate equal to the duplication rate, AD $\approx$ 20%, and $\approx$ 40% MGTE.

| Method | Mean | Median |
|---|---|---|
| Figure 6(a): 1000 genes | | |
| DISCO+QR (LE) | 0.066 | 0.044 |
| STRIDE | 0.054 | 0.030 |
| Figure 6(b): 500 genes | | |
| DISCO+QR (LE) | 0.070 | 0.048 |
| STRIDE | 0.060 | 0.044 |
| Figure 6(c): 100 genes | | |
| DISCO+QR (LE) | 0.081 | 0.075 |
| STRIDE | 0.262 | 0.077 |

Table S7: Means and medians for all model conditions from Figure 6. Error (nCD) of STRIDE and DISCO+QR on a simulated dataset with 1000 taxa (and one outgroup taxon) and varying numbers of genes. (a): 1000 genes, (b): 500 genes, (c): 100 genes. The species tree was estimated with ASTRID-DISCO. The simulated model conditions contain 10 replicates, a duplication rate of $5 \times 10^{-10}$ and an equal loss rate, AD $\approx$ 20%, and $\approx$ 40% MGTE.
